# Nanoscale protein clustering modulates the input-output response of cellular signalling pathways

**DOI:** 10.1101/2025.03.13.642965

**Authors:** Luca Panconi, Juliette Griffié, Dylan M Owen

## Abstract

A key function of the plasma membrane is regulating signal transduction. Proteins involved in the transduction process play a signal-processing role; mapping an input (e.g. number of engaged receptors) to an output (e.g. level of downstream phosphorylation). In many cases, a digital mapping is desirable, i.e. that a cell activates or responds fully once a set input threshold is surpassed. It is believed that the nanoscale organisation of proteins, such as their clustering, modulates this behaviour by altering the frequency of protein-protein interactions. Here, we develop a generalised simulator for dynamic molecular clustering built around agent-based modelling. We show that the clustering properties (e.g. size of clusters, percentage of monomers) tunes the cellular response. This work paves the way for designing potential therapeutic interventions that alter the nanoscale patterning of molecules on the cell surface in order to alter cell behaviour.

**AUTHOR SUMMARY:** Signal transduction is the process by which cells convert external signals into internal responses. This is essential for such functions as hormone signalling and immunity. Many membrane proteins form nanoscale clusters on the cell surface, and state-of-the-art imaging methods allow us to see these clusters in detail. We developed an agent-based simulator to explore how clustering of key signaling proteins (such as kinases and phosphatases) can influence cell signalling outputs. Our results suggest that protein clustering can “digitize” signals, pushing cells to respond in an “all-or-nothing” manner, rather than producing a smooth range of responses. This research contributes to a broader understanding of how dynamic reorganization of transmembrane proteins contributes to information processing. This is a central question in systems biology, with relevance to the fields of immunology, neuroscience, and cancer. By highlighting how the geometric properties of protein distributions can alter signalling responses, we may identify novel methods to manipulate cell behavior.

## INTRODUCTION

Signal transduction across the plasma membrane is vital. This includes, but is not limited to, the function of hormone receptors, ligand-gated ion channels, the T/B receptors of immune cells, the regulation of adhesion molecules and others[1–4]. In all cases, a number of proteins must work in concert to transduce the signal and frequently proteins in the signalling pathway will compete – for example, the opposing functions of kinases and phosphatases on a particular target[5, 6, 7]. An under-explored aspect of these functions is to consider the input-output, or signal-processing response of a pathway[8]. The input is the strength of the signal initiating transduction. This can be variously interpreted, but might for example be the number of engaged receptors, or the amount of time that those receptors have been engaged[9, 10]. The output is then the downstream response. Again, there may be many definitions, but it might include the phosphorylation level of some downstream master regulator, or even changes in gene expression[11]. In general, any relationship between input and output may be possible, but it will be broadly determined by the many proteins that make up the pathway and how they interact including, for example, positive and negative feedback[12].

It is now widely regarded that many membrane and membrane-associated proteins do not display random distributions on the cell surface. Instead, they show a rich variety of organisations including dimerization, nanoscale clustering and large, microscale aggregates[13, 14, 15]. These behaviours are observed despite almost all proteins remaining mobile on the surface with a range of (usually non-Brownian) diffusive behaviours[16, 17]. The non-random distributions are usually attributed to a range of biophysical phenomena associated with the membrane including, but non limited to, direct protein-protein interactions, membrane lipid microdomains, corralling by the cortical actin cytoskeleton and active processes like endo- or exocytosis[18–20]. The nanoscale clustering of proteins is now a measurable phenomenon owing to the development of electron microscopy and, more-recently, super-resolution fluorescence imaging such as single-molecule localisation microscopy (SMLM)[21]. Combined with statistical analysis, these allow the quantification of clustering properties such as cluster sizes, monomer fractions, cluster density and so on[22–25]. The motivation for such studies is frequently that the nanoscale distribution of proteins will modulate the frequency of their interactions (collisions) and therefore can influence the behaviour of the aforementioned input-output signal processing[3, 11]. Despite this, how a particular nanoscale distribution of a protein influences the behaviour of the signalling pathway(s) it is a member of remains poorly understood. Due to experimental limitations (e.g. tools for manipulating nanoscale organisation), the relationship between protein distribution and signalling pathways behaviour is best studied computationally. Previous studies for example, have shown that the presence of protein clusters can digitise cellular signalling[6].

Here, we present and demonstrate a simulation ecosystem built around agent-based modelling (ABM)[16]. The simulator can generate specified clustered distributions in which all molecules remain diffusive and dynamic. By incorporating multiple protein species (e.g. a target together with a competing kinase and phosphatase), we can use the simulator to test how nanoscale organisation influences the input-output behaviour of a simple signalling pathway. In this case, we take the input to be the ratio of activator (the kinase) to inhibitor (the phosphatase) and the output to be the fraction of phosphorylated receptors. We show that clustering of the pathways constituent components does lead to digitisation of the fraction of simulations (cells) that pass a specified output threshold. We further show how the exact clustering properties (such as cluster size) influence the shape of this relationship.

## Results

### Agent-based modelling as a method for simulating protein aggregation

In this work, we build upon ABM approaches and use Ripley’s K-function as a metric to direct the evolution of a point cloud from an initial condition where points are randomly distributed towards a target distribution. The algorithm can be initialised with a (simulated or experimentally measured) target point pattern or a (simulated or experimentally derived) target K-function (see **Supplementary Information**). Through an iterative Markov process, each point is sequentially transposed with step size proportional to the error between its own K-function and the global target. As the individual error of each point is reduced, so too is the step size, which promotes aggregation at scales summarised by the target K-function (**Figure 1a**). This culminates in a model which minimises the error between the measured global K-function of a point pattern and the target. The process is designed to simulate point clouds with defined clustering statistics while having all points remain mobile, even after convergence of the measured and target functions (**Figure 1b**).

**Figure 1:**
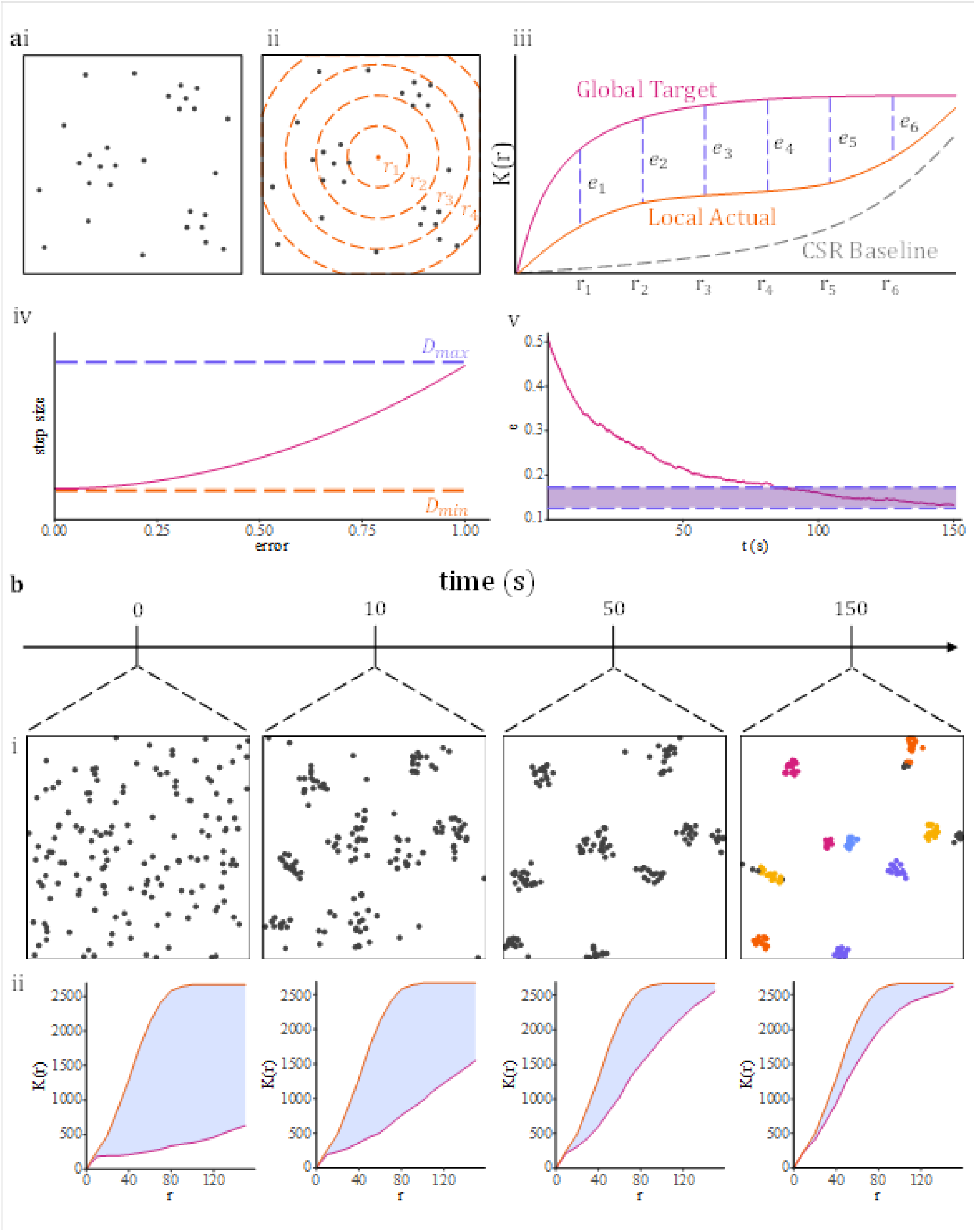
**a** An example target point cloud (i) is analysed (ii) to generate a global, target Ripley’s K-function (iii). From this the error (e) between the cloud’s actual K-function and the target is calculated and averaged at each radius. This error is then mapped to a step size (iv) between minimum and maximum values (𝐷_𝑚𝑖𝑛_ and 𝐷_𝑚𝑎𝑥_) via a quadratic function. Over many iterations, this causes the actual K-function to converge to the target (v) defined at the point where the variance of the error falls below 0.05. **b** Example of the evolution of a point cloud under this framework (i) and the corresponding Ripley’s K-functions (ii) of the simulation (magenta) and target (orange).

To test the efficacy of our model, we simulated a series of target point clouds with varied cluster properties over 2µm × 2µm regions of interest (ROIs). We used a maximum diffusion coefficient of 𝐷 ∼ 0.1𝜇𝑚𝑠^−2^, which equates to a step size of 𝛥𝑥 ≈ 63𝑛𝑚 per 10𝑚𝑠 time frame (see **Materials and Methods**). After the model reaches convergence, we then perform DBSCAN cluster analysis to extract cluster descriptors and compare them to the original target parameters[27]. For each simulation, we tracked the point pattern distribution and the measured K-function relative to the target. Further, we record the difference between the measured number of clusters, the measured number of points per cluster, and their corresponding targets over time (**Figure 2a**). Both the number of clusters and points per cluster converged to within 10% of their input target values.

**Figure 2:**
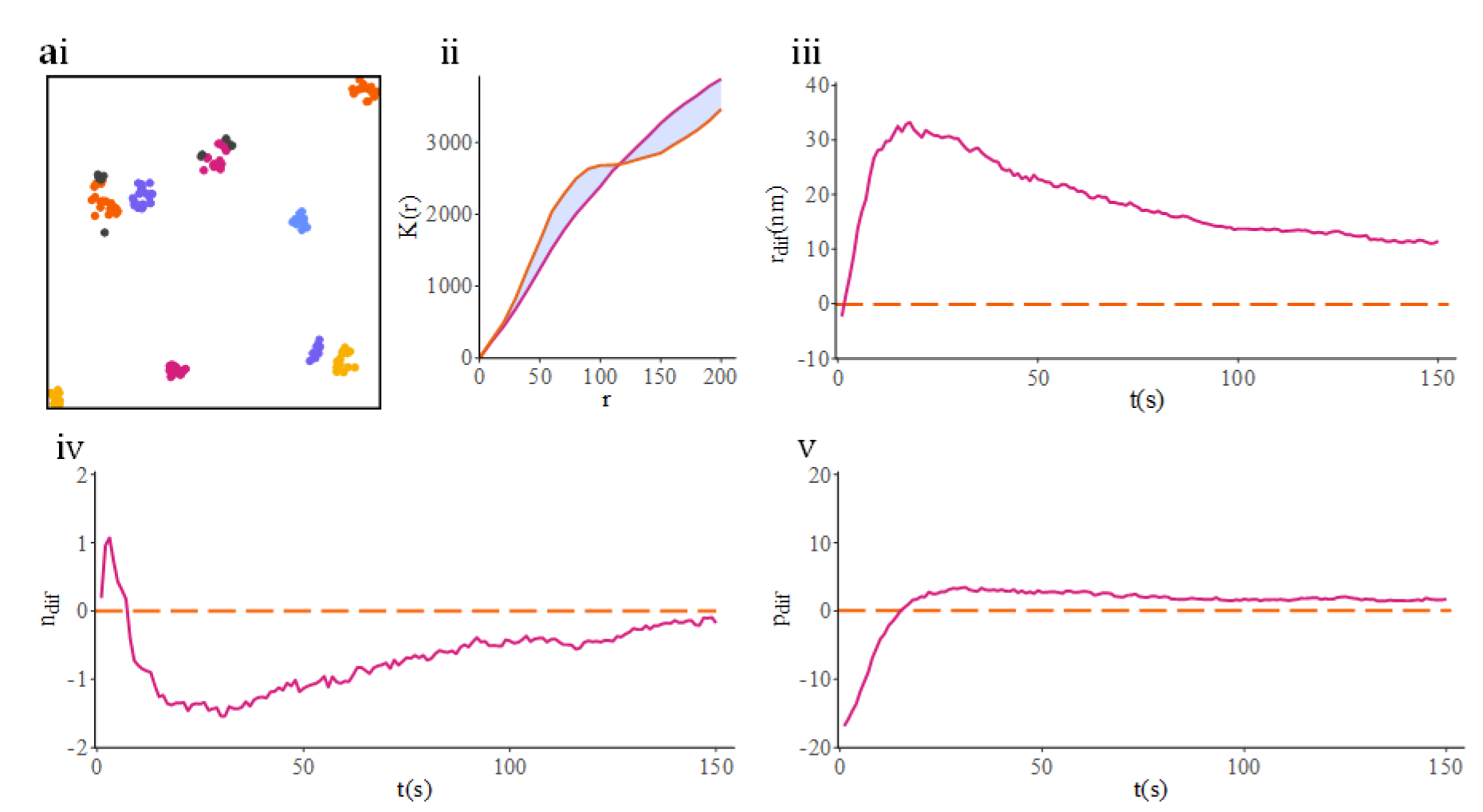
Convergence of cluster properties in simulated and experimental data. **a** Simulated data. (i) Coloured points represent clusters identified by DBSCAN. (ii) The difference between the Ripley’s K-function (orange) and the target function (magenta). Average difference between simulation and target (iii) cluster radius (r_dif_), (iv) number of clusters and (v) number of points per cluster.

### Modelling interactions between multiple molecular species

Extending upon our single-population model, we have developed a system for simulating multiple interacting molecular species (see **Materials and Methods**). Each population may be aggregated separately or co-clustered. We consider an arbitrary three-population model, including activators and inhibitors and their receptor targets (which may be active or inactive). We tested each of the 15 geometric configurations given in **Figure 3a-o** and described in **Table 1**. Receptors are activated or inhibited within a specified range (5nm) of activators or inhibitors, respectively, and remain in their state until acted upon again (**Figure 3p**). For each configuration, we considered 64 possible ratios of activator and inhibitor populations and undertook 20 trial simulations for each. This experiment was repeated 9 times, giving a total of 11520 simulations per configuration, with each simulation lasting 2.5 minutes. The percentage of receptors which were active at system convergence was recorded (**Figure 3q**). The probability of cell activation for each A/I ratio was estimated as the percentage of simulations in which the proportion of activated receptors exceeded 0.5 (**Figure 3r**). A transition function was estimated by fitting a sigmoidal curve (**Figure 3s**). From this curve, two parameters were estimated: 𝛼, the point of inflection, and 𝛽, the transition rate or scale (see **Supplementary Information** for formula). Here, 𝛼 may be interpreted as the A/I ratio above which the system is likely to induce cell activation. The lower the value of 𝛽, the greater the increase in probability of cell activation with respect to the increase in A/I ratio.

**Figure 3:**
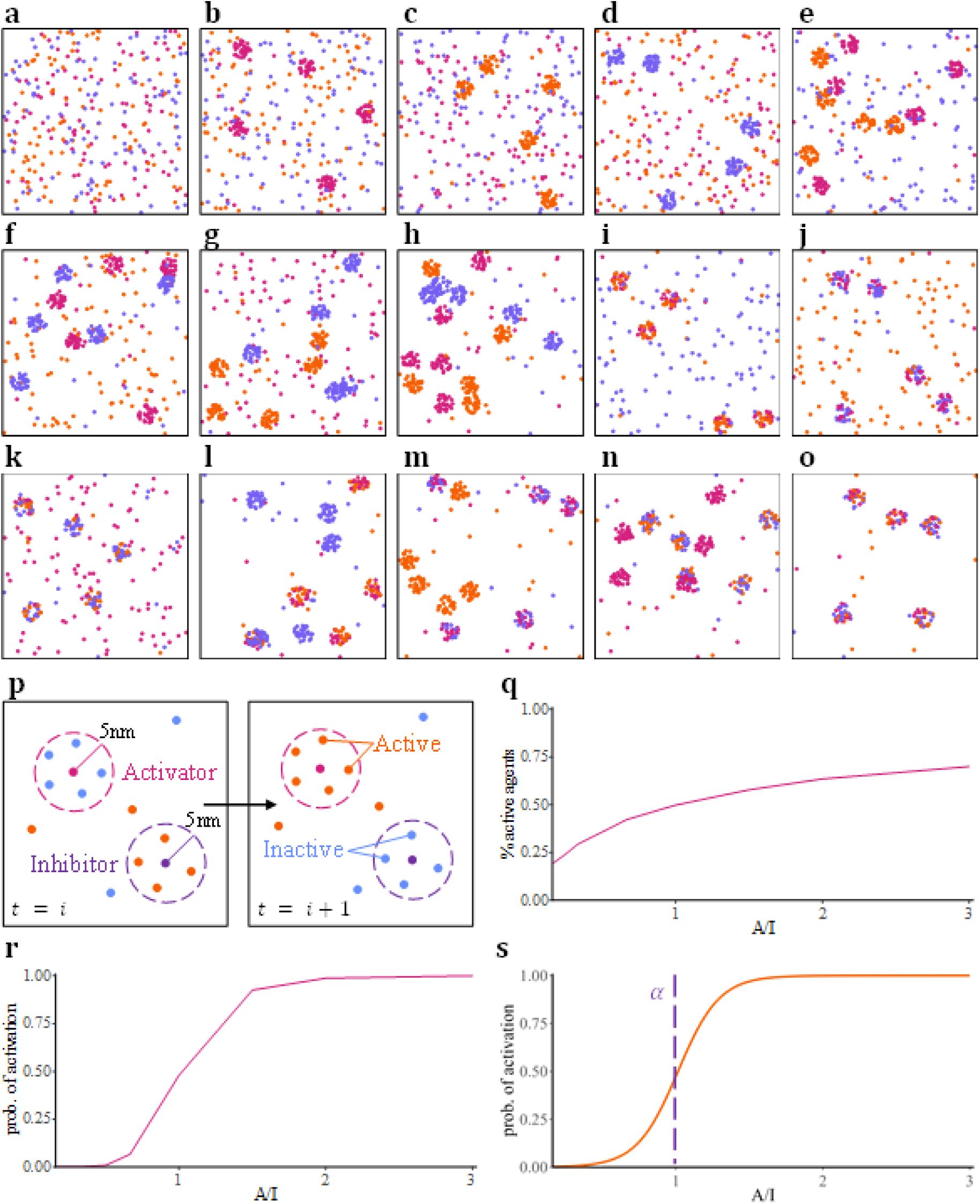
Modelling multiple populations. **a-o** Three interacting molecular species under varied geometric point pattern configurations. **p** Receptors (orange) become active when in a 5nm radius of an activator (magenta) and remain active until coming within a 5nm radius of an inhibitor (purple). **q** The percentage of active receptors at system convergence versus activator-inhibitor ratios (A/I) for the case for all distributions being CSR. **r** Estimated probability of cell activation (% of simulations in which the % of activated receptors exceeds 50%). **s** Fitting a sigmoidal growth function to the plot in **r** and estimating transition parameters. 𝛼: point of inflection, 𝛽: transition rate (scale). Here, 𝛼 = 1.01 (purple dashed line) and 𝛽 = 0.14.

**Table 1:** Description of labels for each geometric point pattern configuration given in Figure 3.

| Label | Configuration Description |
| --- | --- |
| **a** | All populations at CSR. |
| **b** | Clustered activators, with all other populations as CSR. |
| **c** | Clustered receptors, with all other populations as CSR. |
| **d** | Clustered inhibitors, with all other populations as CSR. |
| **e** | CSR inhibitors, with all other populations clustered. |
| **f** | CSR receptors, with all other populations clustered. |
| **g** | CSR activators, with all other populations clustered. |
| **h** | All populations clustered. |
| **i** | Co-clustered activators and receptors, with inhibitors as CSR. |
| **j** | Co-clustered activators and inhibitors, with receptors as CSR. |
| **k** | Co-clustered receptors and inhibitors, with activators as CSR. |
| **l** | All populations clustered, with activators and receptors co-clustered. |
| **m** | All populations clustered, with activators and inhibitors co-clustered. |
| **n** | All populations clustered, with receptors and inhibitors co-clustered. |
| **o** | All populations co-clustered. |

Points of inflection (**Figure 4a**) and scales (**Figure 4b**) were recorded for each configuration, over all 9 trials. Boxplots depicting the distribution of estimated parameter values are given in **Figure 4ai** and **4bi**. An ANOVA test for evidence of difference between configurations returned a significant p-value for both parameters, suggesting statistically significant evidence of differences in points of inflection and scales between configurations. A Tukey Honest Squares Difference (HSD) test was performed for both parameters, across all configurations. Results for differences in point of inflection and scale are given in **Figure 4aii** and **4bii** respectively. We observe the lowest inflection point (i.e. minimum A/I ratio required to induce activation) in configurations **i** and **l**, when activators and receptors are co-clustered, and highest inflection point when receptors and inhibitors are co-clustered.

**Figure 4:**
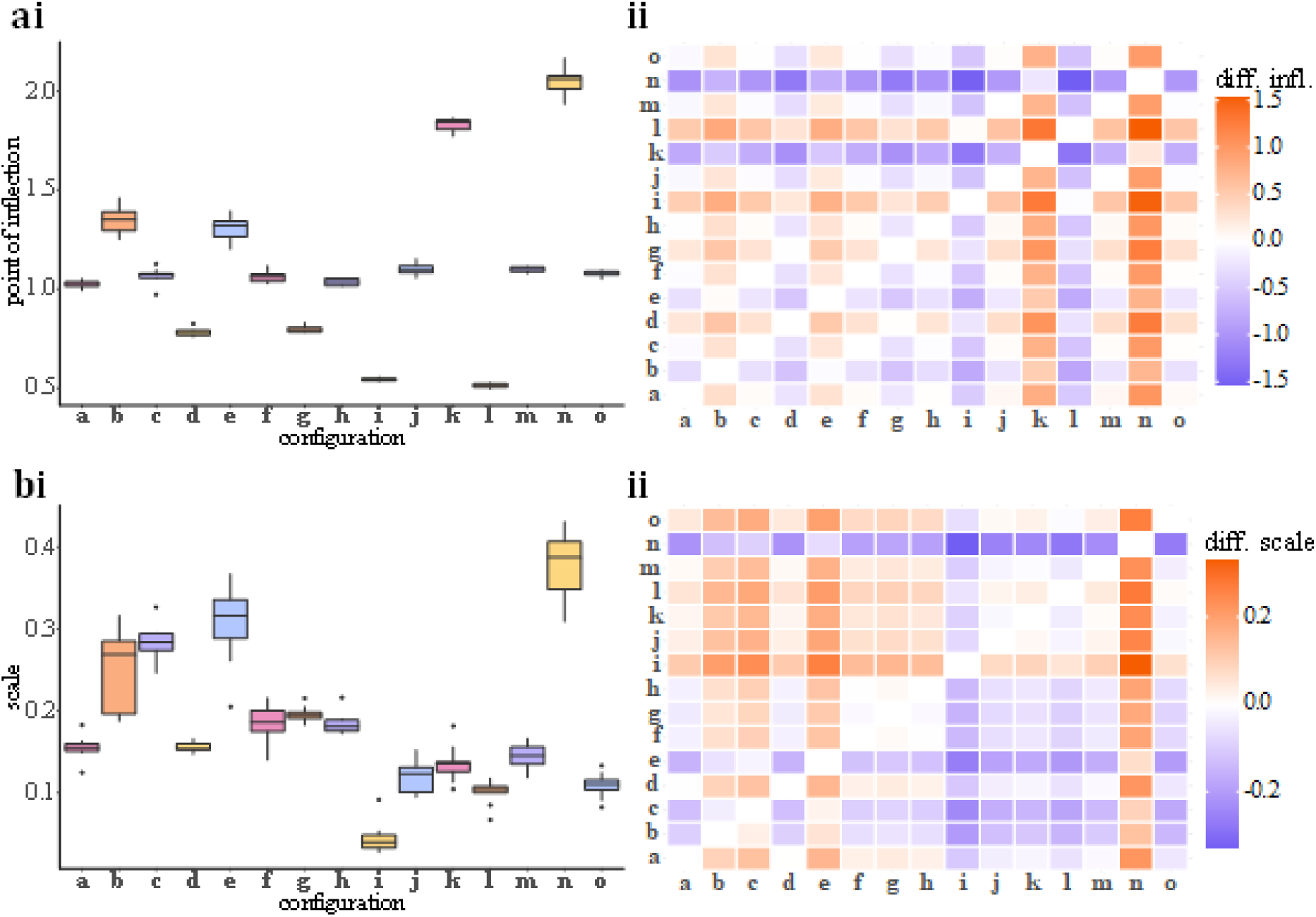
Results of three-population simulations under varied configurations (as shown in Figure 3). **a** (i) Boxplots of all points of inflection estimated from sigmoidal fits. (ii) Differences in point of inflection for each pair of geometric configurations, as returned by Tukey’s Honest Significant Difference (HSD) test. **b** (i) Boxplots of all scale parameters estimated from sigmoidal fits. (ii) Differences in scale parameters for each pair of geometric configurations, as returned by Tukey HSD test.

### Receptor cluster properties regulate signal propagation

To measure the contributions of each cluster property towards the induction of cell activation, we again consider three-population simulations. Here, we record the estimated probability of cell activation while varying individual cluster properties. For this particular use case, we consider only clustered activators, while all other distributions are taken as CSR. We show the impact of perturbing cluster radius (**Figure 5a**), number of clusters (**Figure 5b**) and the percentage of points (proteins) assigned to clusters (**Figure 5c**) for a range of A/I ratios. Outputs for the full range of A/I ratios considered in these simulations are given in **Supplementary Figures 1-3**.

**Figure 5:**
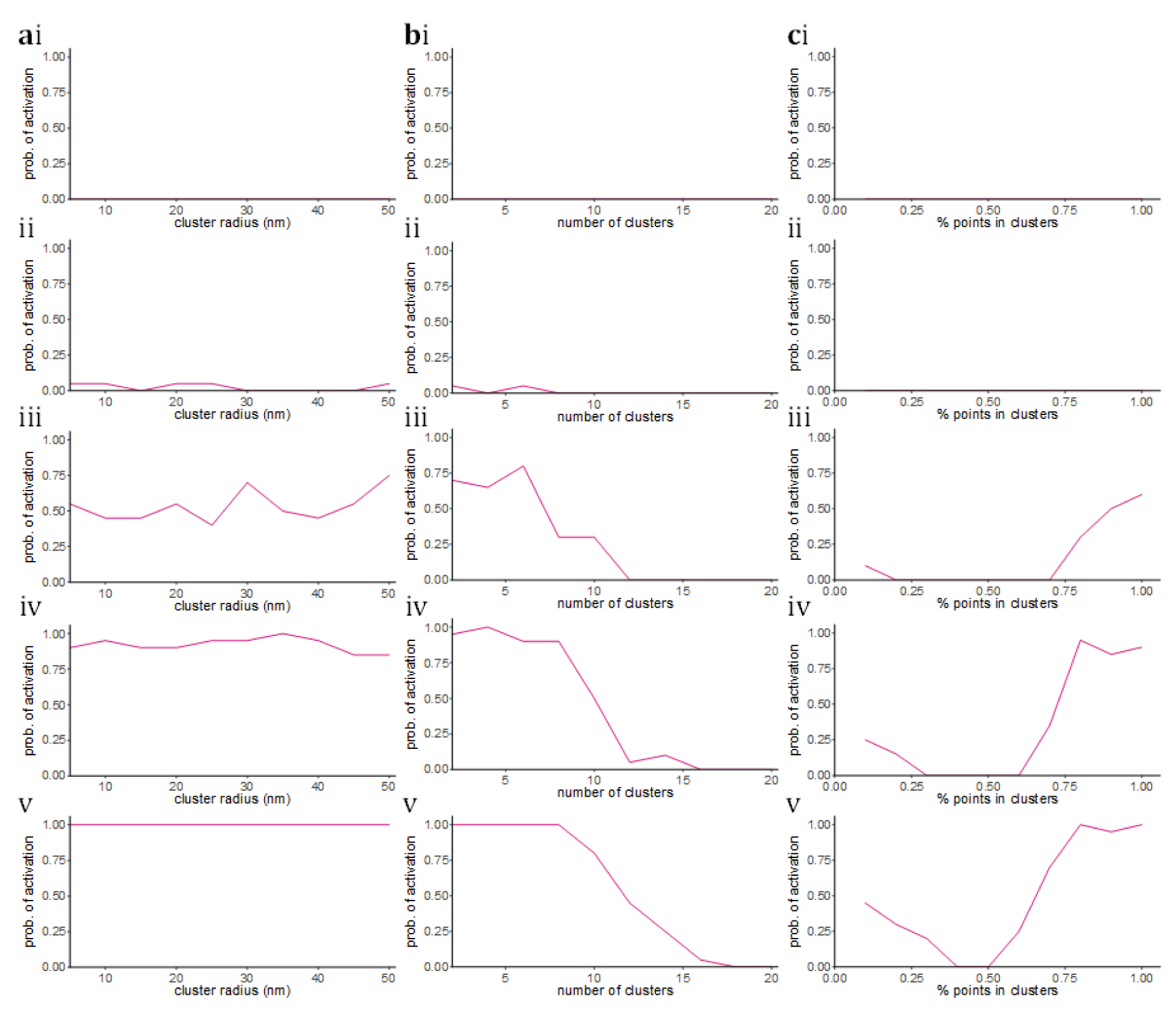
Estimated probability of cell activation versus individual cluster properties in the case where activators were clustered, while all other distributions were CSR. **a** Cluster radius versus probability of cell activation. **b** Number of clusters versus probability of cell activation. **c** % points assigned to clusters versus probability of cell activation.

Results suggest that the probability of cell activation remains approximately constant for cluster radii between 5nm and 50nm. Furthermore, increasing the number of clusters generally decreases the probability of activation. This may suggest that activation favours fewer, denser clusters, as opposed to a greater number of sparse clusters. The activation probability also shows a highly non-linear relationship against the percentage of points assigned to clusters. This may suggest that activation probability can be minimised by a trade-off between the percentage of activators within the CSR background and the percentage in clusters. In this case, simulations with a higher fraction of monomers allow the activators to move faster, making them more likely to interact with receptors. This interpretation is validated in **Supplementary Figure 4**, in which we record the average step size (per simulation) versus the percentage of points per cluster and show a decrease in step size as the percentage of points in clusters increases.

## Discussion

Membrane protein aggregation underpins essential cellular processes such as signalling and communication. Central to this notion is the spatial clustering of membrane receptors, which is hypothesised to convert analogue extracellular signals into digital intracellular counterparts. In this work, we have developed an MCMC simulation method to model this process. While the framework is generalizable, here we investigate the simple system of activators (e.g. a kinase) and inhibitors (e.g. a phosphatase) acting on the same target (e.g. a receptor). The simulator allows us to specify a desired cluster configuration for all three molecule types, monitor the receptor activation (phosphorylation) level and then analyse the fraction of simulations (cells) that surpass a specified activation threshold. Many transmembrane proteins form nanoscale clusters, and while it was hypothesized that these clusters convert analog extracellular signals into digital intracellular signals, the mechanism of digitization is not fully understood[6]. Here, we have shown that an increase in activator-inhibitor ratio produces a transition function in the form of a sigmoid activation curve, whose parameters vary depending on the cluster configuration being modelled.

Our results show that the clustering properties of the molecules profoundly influence the probability of activation. In particular, the co-clustering of activators and receptors increases the activation probability. Likewise, the co-clustering of inhibitors and receptors decreases the activation probability. The exact level of activation can be fine-tuned by altering the clustering properties (e.g. cluster sizes, percentage monomers and so on). This platform provides a versatile system for testing, evaluating and comparing hypotheses surrounding dynamic reorganization of transmembrane proteins. In particular, PAD simulations could be used to estimate the geometric properties of phosphatase/kinase distributions which maximise the probability of inducing signal propagation. By modelling receptor distributions under treatment with cross-linking agents or nanotherapeutics (which may be determined from molecular localization data), it is possible to estimate signalling behaviour without further *in vitro* experiments. This could be used to inform the design of nano-medicines which explicitly re-organise transmembrane receptors to achieve signal activation (e.g. coated DNA-Origami flat sheets[28]), therefore opening the possibility of developing therapeutic agents which model protein clustering in a controlled way.

## Materials and Methods

### PAD simulation software

The ASMODEUS software package (v. 1.0.0) was written in the R programming language (v. 4.4.2) and employed in the integrated development environment RStudio, (2022.07.1+554). ASMODEUS is available for use under GNU General Public License (v. 3.0).

### Simulating target distributions

Target distributions were simulated by generating a fixed number of circular clusters with specified cluster radius and number of points per cluster, and then overlaying outliers subject to a given background ratio. Cluster centre coordinates were selected randomly such that all clusters would be contained within the ROI. Cluster points were uniformly generated around centres at distances of 𝑟_𝑡_𝑋^2^, where 𝑋∼𝑈𝑛𝑖𝑓(0,1) and 𝑟_𝑡_is the target cluster radius. The range of parameter values used in single-population simulations (unless otherwise specified) is given in **Table 2**.

**Table 2:** Parameters used to simulate target distributions.

| Parameter | Function | Values |
| --- | --- | --- |
| $n_t$ | Number of clusters | 10 |
| $r_t$ | Cluster radius. | 10nm, 30nm, 50nm |
| $p_t$ | Number of points per cluster. | 5, 15, 30 |
| $b_t$ | Background to cluster ratio. | 0, 0.33, 0.66 |

### General PAD simulations

First, 100 simulations were run over 100,000 frames and the maximum time taken to achieve convergence was measured at 7529 frames. As such, the maximum frame number for all future simulations was taken at double this value, rounded to the nearest 1000, to increase the probability of convergence. Therefore, unless otherwise specified, each simulation was run over 15000 frames, with each frame representing 10ms of real time, yielding a total simulated time of 2.5 minutes. All simulations converged within the allocated time. The domain size was fixed at 2µm × 2µm, a standard ROI size for super-resolution microscopy. A diffusion coefficient of 𝐷 ≈ 0.1μm^2^/s was selected – this represents the diffusion coefficient of a T cell receptor (0.082μm^2^/s, rounded to the nearest 10^−1^), taken as a model molecular species[17]. Note that this framework could be applied to any protein with known diffusivity coefficient and TCR serves only as an exemplar here. This equates to a step size of 𝛥𝑥 ≈ 63𝑛𝑚 per 10𝑚𝑠 time frame (see **Supplementary Information**), and so 𝐷_𝑚𝑎𝑥_ = 63𝑛𝑚 was set[26]. The average error is recorded at each frame. We defined the convergence time as the first frame in which the variance in the error of all future frames fell below 0.05. When applicable, DBSCAN cluster analysis[29] was performed to extract cluster descriptors, for comparison with the original target parameters.

### Simulating multiple populations

Simulations can incorporate an arbitrary number of distinct molecular species simulated within the same ROI. Species can be simulated separately, by giving each population a separate target and ignoring species outside their own, or co-clustered with other populations, by giving a subset of the populations the same target and only considering the density of species within that subset. Interactions between populations can be recorded and tracked across consecutive frames. In this work, we consider a three-population model, including activators, inhibitors and receptors (which may be active or inactive), depending on interactions with the former two molecular species. In particular, receptors are activated within 5nm of activators and inhibited within 5nm of inhibitors. All receptors are initialised as inactive. By outputting the resulting point pattern of any given simulation into another, any of the above regimes can be modulated and concatenated.

## Author Contributions

LP: Conceptualisation, data curation, formal analysis, funding acquisition, investigation, methodology, project administration, resources, software, validation, visualisation, writing - original draft preparation, writing - review and editing. JG: Conceptualisation, funding acquisition, methodology, project administration, supervision, writing - review and editing. DMO: Conceptualisation, funding acquisition, methodology, project administration, supervision, writing - review and editing.

## Conflict of Interest

The authors declare no financial or commercial conflict of interest.

## Data Availability Statement

The software package and data that support the findings of this study are available at: https://github.com/lucapanconi/asmodeus. ASMODEUS v. 1.0.0[30] and associated data are available for public use under the terms of the GNU General Public License v. 3.0 (DOI: https://doi.org/10.5281/zenodo.12627125).

## Disclosure of funding sources

JG acknowledges funding from the Knut and Alice Wallenberg Foundation Data Driven Life Sciences grant (31003604). LP acknowledges funding from the SciLifeLab RED Postdoctoral Fellowship (31005394). Dylan Owen acknowledges funding from the Biotechnology and Biological Sciences Research Council (BBSRC), grant BB/X018644/1.

## Supplementary Information

### Simulating PAD through MCMC agent-based modelling

Our model is defined under the assumption that proteins are represented by infinitesimal points, which move laterally across the plasma membrane, approximated by a 2D surface. Protein movement is stochastic, with agents moving in a random direction subject to a deterministic step size. This step size, in turn, is controlled by a spatially descriptive statistic (here, the Ripley’s function), derived from the target point pattern. The algorithm for simulating the model is as follows. First, input a target point pattern and appropriate simulation parameters: this includes an expected maximum step size, Dmax, an ROI size, and a time frame over which to run the simulation (for guidance on parameter selection, see below). Calculate the Ripley’s K-function from the target distribution and initialise a completely spatially random (CSR) distribution. For each point, calculate the error between the localised K-function and global target K-function, and then determine the step size (for formulas, see below). Offset each point in a randomly selected direction, apply edge correction to return all points to the domain. A schematic of the algorithm (**Schematic 1**) and pseudocode are given below.

### Parameter selection

Large scale emergent behaviour in ABMs is population sensitive, so the number of agents must be able to reflect the reality of the system being modelled. The precise number of agents found in the target distribution is used automatically. It is generally straightforward to determine the point pattern of protein localisations from raw super-resolution data, from which the molecular counts, ROI size and corresponding spatial statistics are automatically determined in our pipeline. All that remains is to specify appropriate minimum and maximum molecular velocities, proportional to the diffusion coefficient, which can be derived experimentally or mathematically. For applications to experimental data, expected diffusivities can be quantified from imaging methods such as single particle tracking or fluorescence correlation spectroscopy. The expected step size 𝑥 taken in time 𝑡 is then estimated by 𝑥 = √2𝐷𝑡, where 𝐷 is the diffusion coefficient.

### Limitations of model assumptions and proposed solutions

The foremost limiting assumptions of our model are as follows:

- Proteins are represented by infinitesimal points and therefore do not collide. This can be circumvented by rejecting protein movements if they come within a range (equal to the expected radius of the molecule) of other proteins and either a) redistributing or b) reflecting the proteins at the angle of incidence, with distance equal to the step size not yet covered.
- The plasma membrane is approximated by a 2D surface and proteins move laterally in the plane of the membrane. However, the Ripley’s function, as a spatially descriptive statistic, is based only on distance and density and can therefore be applied to any number of spatial dimensions. This, with correction for topographic variation, helps circumvent clustering artefacts that could arise from membrane undulations and curvature projected into 2D data^28,71^.

### Calculation of the Ripley’s functions

The localised Ripley’s K-function for a point 𝑖 is given by,

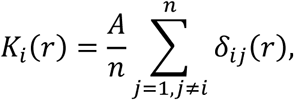

where 𝐴 is the area of the ROI, 𝑛 is the total number of points, 𝑟 is the search radius and 𝛿_𝑖𝑗_(𝑟) is the indicator variable that equals 1 if point 𝑗 is within a radius 𝑟 of point 𝑖 (and 0 otherwise). This represents the spatially-averaged density localised around point 𝑖. Averaging over all points gives the global Ripley’s K-function,

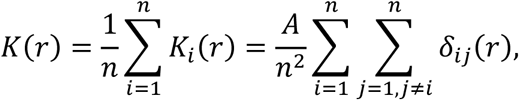

which represents the spatially averaged density across the ROI. The target function derived from the input distribution is denoted 𝐾_𝑡_(𝑟).

### Derivation of error and step size

The average ratiometric error of point 𝑖 is denoted by,

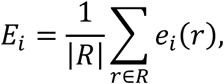

where 𝑅 is the finite set of radial values to iterate over (derived during calculation of the Ripley’s function) and 𝑒_𝑖_(𝑟) is the scalar error at radius 𝑟 given by,

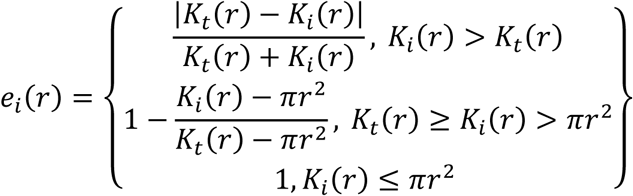

The lower bound is set at 𝜋𝑟^2^as this is the theoretical form of the K-function for a CSR distribution. The set 𝑅 is granularised for ease of computation and to prevent overfitting. The step size of point 𝑖 is then given by,

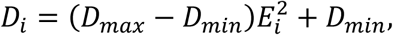

where 𝐷_𝑚𝑖𝑛_is an optional minimum step size parameter, which defaults to 0. A quadratically decreasing step size was chosen based on previous results^34^.

### Formula for sigmoidal transition function

The formula for a sigmoidal curve is given by,

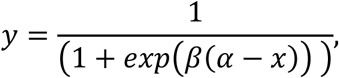

where 𝛼 is the point of inflection and 𝛽 is the scale parameter.

**ASMODEUS Pseudocode**
#Initialise:
#Dmin - Minimum step size.
#Dmax - Maximum step size.
#ROI - A vector of two values representing the width and height of the region of interest, respectively. The region of interest is considered from the origin, so pointCloud coordinates must be translated accordingly.
#times - Number of time steps or iterations of the simulation.
#pointCloud - A matrix or data frame with two columns: x and y coordinates of each point. Used as a target distribution from which target K function is drawn.
#rmax - Numeric value, the maximum radius for each K function to be calculated at.
#nrval - Numeric value, the number of equally-spaced radial values for each K function to be calculated over.
#target - Manual input for target function. Can be left as null as long as a target pointCloud is provided.
#numberOfPoints - Number of points to use in simulation.
#initialDistribution - The initial frame of the simulation, used to specify pre-determined spatial organisation. If left null, a uniform random distribution will be used.
**INPUT** Dmin, Dmax, ROI, times, pointCloud, rmax, nrval, target, numberOfPoints, initialDistribution
#Calculate target K function from pointCloud (if pointCloud is given).
**IF** pointCloud is given (not null)

**CALL** RipleyKFunction on *pointCloud*, with r-axis defined on the range of 0 to *rmax* spaced by *nrval* intervals (see **Introduction** for formula)
**SET** output as *target*
**SET** *numberOfPoints* as number of rows in *pointCloud*
**IF** *target* is not given (null)

**END**: not enough information to perform simulation
#Generate initial distribution.
**IF** *initialDistribution* is given (not null)

**SET** currentCloud as initialDistribution
**SET** numberOfPoints as number of rows in initialDistribution
**ELSE**

**SET** *currentCloud* as a completely spatially random distribution, defined over *ROI* with *numberOfPoints* points
#Initialise list of frames to store the point pattern at each time frame.
**INITIALISE** empty list *frames*
#Iterate over each time frame.
**FOR** each time *t* in *times*

#Calculate the K function of currentCloud.
**CALL** RipleyKFunction on *currentCloud*, with r-axis defined on the range of 0 to *rmax* spaced by *nrval* intervals (see **Introduction** for formula)
**SET** output as *currentKFunction*
#Calculate error between currentKFunction and target for each point.
#currentErrors is a list containing the average error for each point in currentCloud.
**CALL** errorFunction on *currentKFunction* to calculate error between *currentKFunction* and *target* for each point (see **Appendix** for formula)
**SET** output as *currentErrors*
#Offset position of each point in currentCloud.
**FOR** each point in *currentCloud*

#Calculate offset for each point using stepSize.
**GET** error *e* of point from *currentErrors*
**CALL** stepSize on *e* with parameters *Dmin* and *Dmax* (see **Appendix** for formula)
**SET** output as *step*
**SET** *randomAngle* to be a random number between 0 and 2 * pi
**SET** coordinates of the point in *currentCloud* by moving point along a path of length *step* in the direction of *randomAngle*
#Once all points have been updated, append this new cloud to the list of frames.
**APPEND** currentCloud to frames
#At the end of this process, frames contains a point cloud for each time frame.
**RETURN** *frames*

**Schematic 1:**
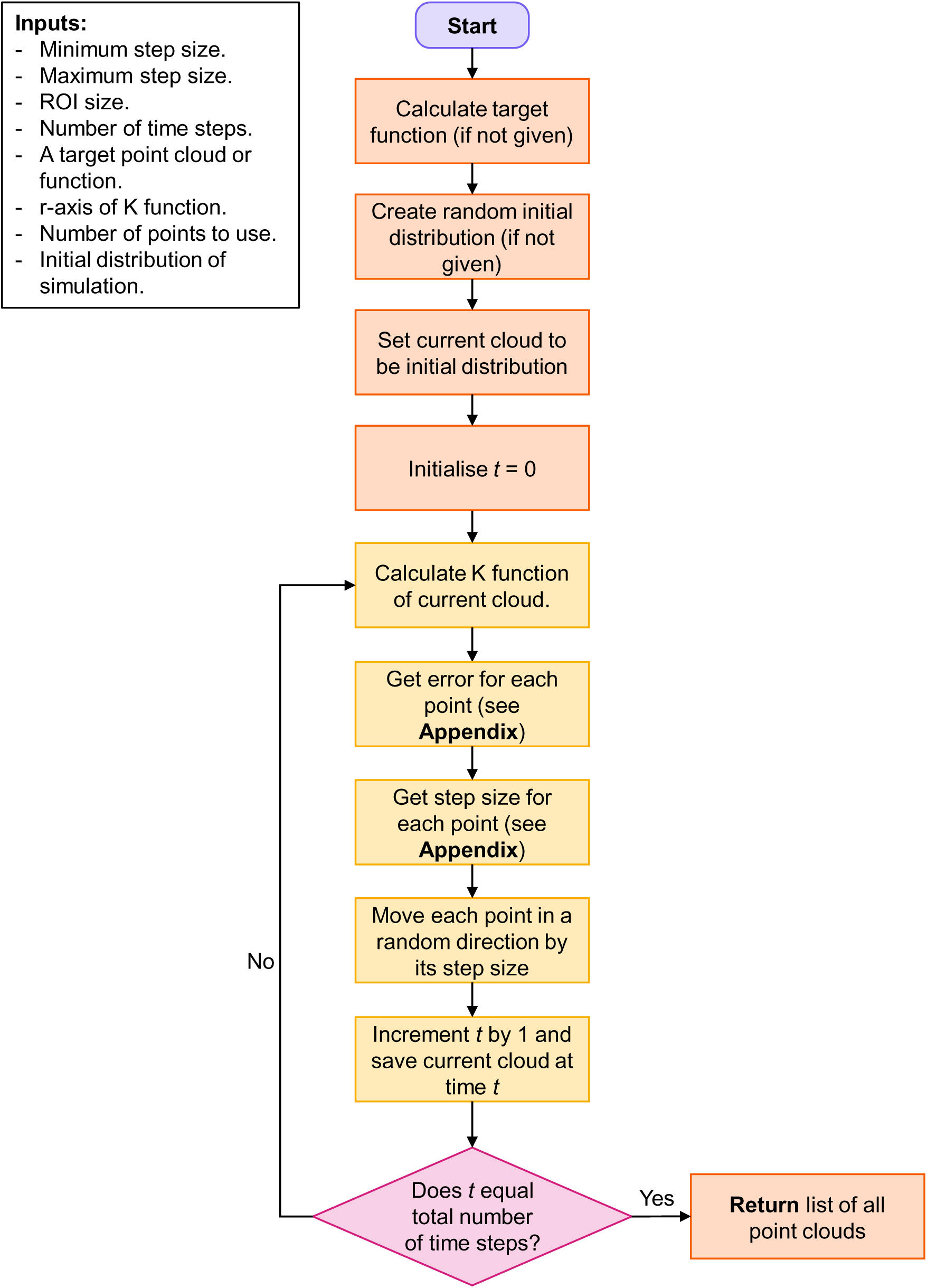
Conceptual diagram of ASMODEUS algorithm.

**Supplementary Figure 1:**
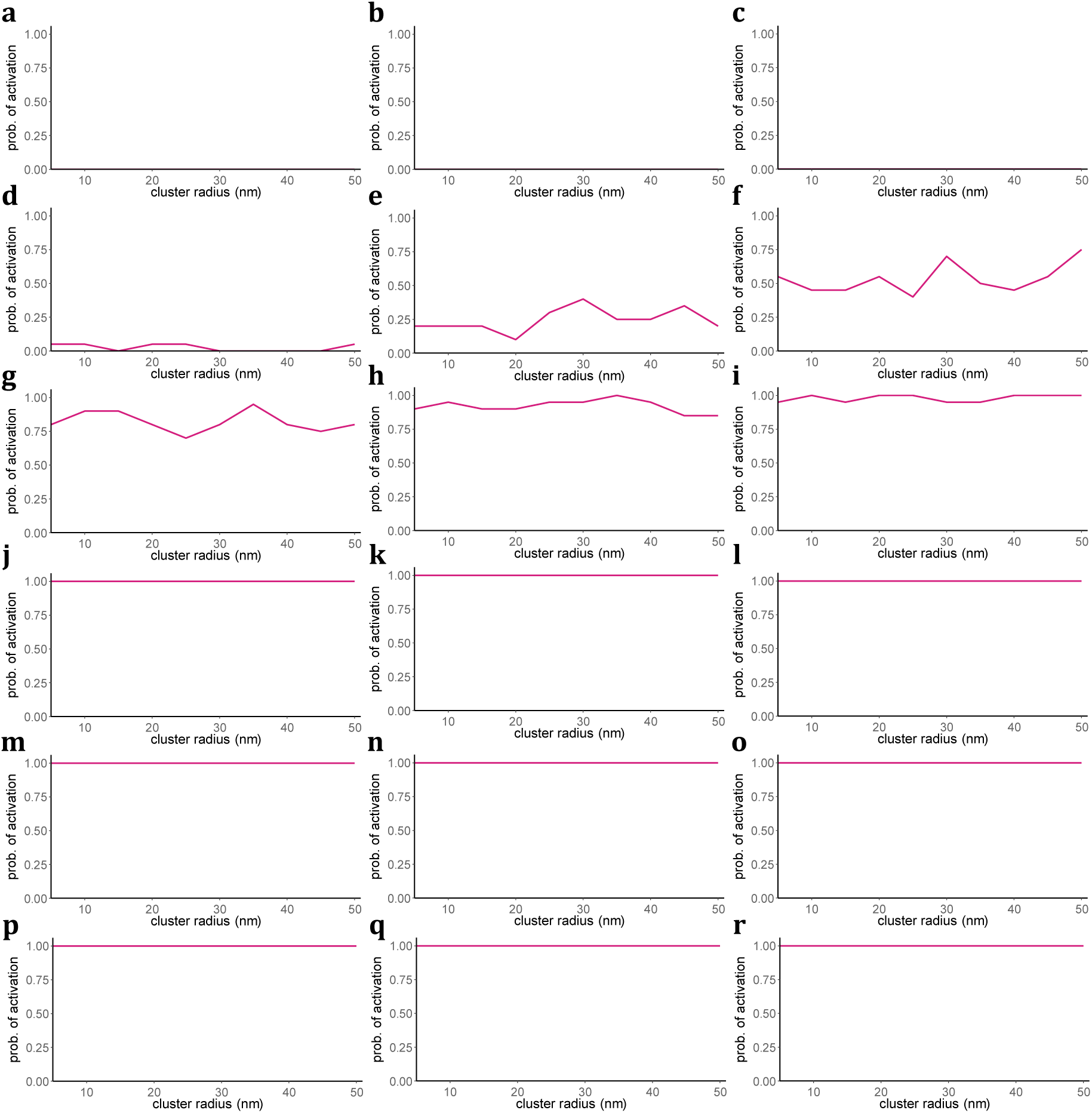
Estimated probability of cell activation versus cluster radius (with all other parameters fixed) for A/I ratios of **a** 0.25, **b** 0.5, **c** 0.75, **d** 1.0, **e** 1.25, **f** 1.5, **g** 1.75, **h** 2.0, **i** 2.25, **j** 2.5, **k** 2.75, **l** 3.0, **m** 3.25, **n** 3.5, **o** 3.75, **p** 4.0, **q** 4.25 and **r** 4.5.

**Supplementary Figure 2:**
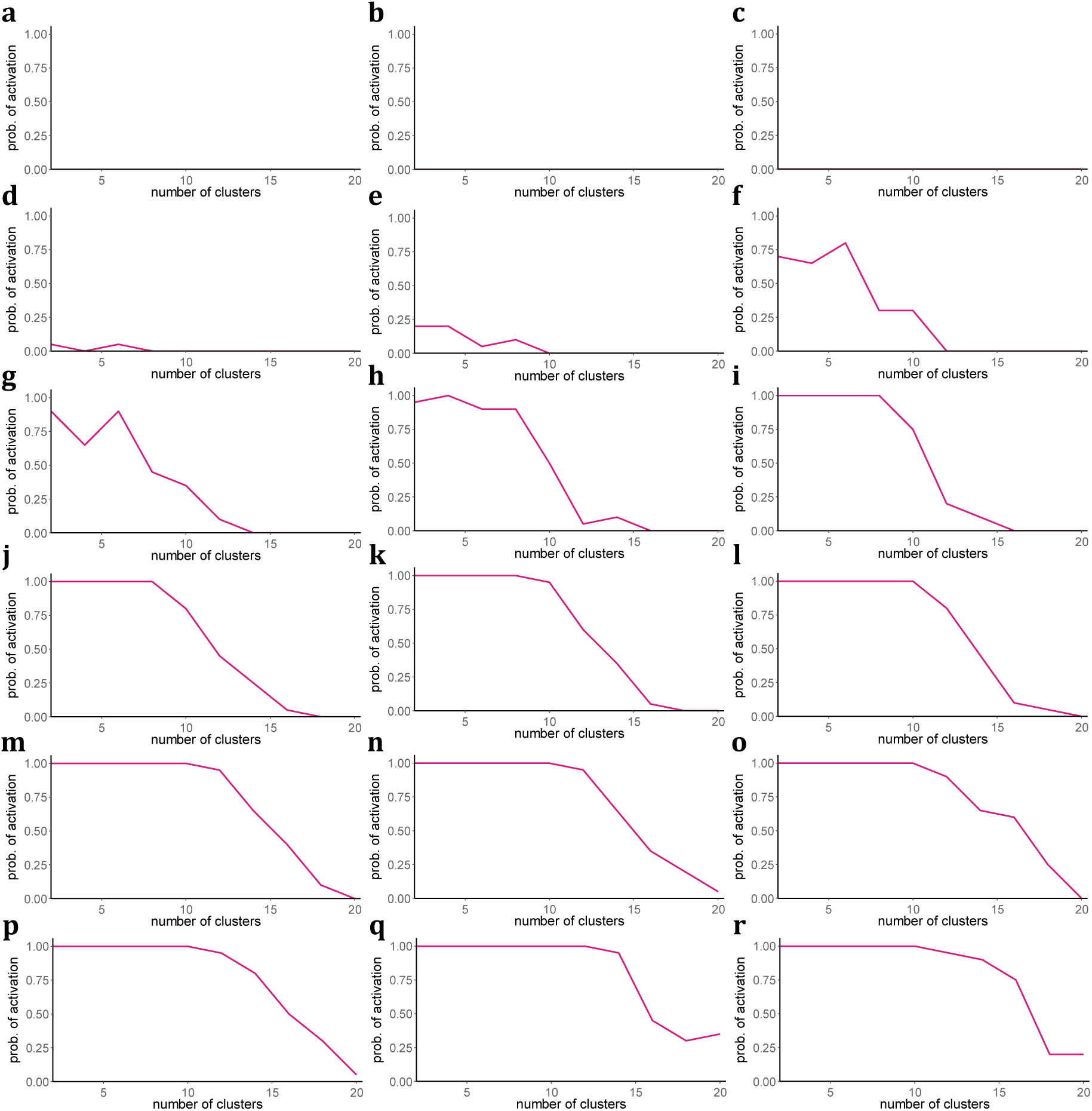
Estimated probability of cell activation versus number of clusters (with all other parameters fixed) for A/I ratios of **a** 0.25, **b** 0.5, **c** 0.75, **d** 1.0, **e** 1.25, **f** 1.5, **g** 1.75, **h** 2.0, **i** 2.25, **j** 2.5, **k** 2.75, **l** 3.0, **m** 3.25, **n** 3.5, **o** 3.75, **p** 4.0, **q** 4.25 and **r** 4.5.

**Supplementary Figure 3:**
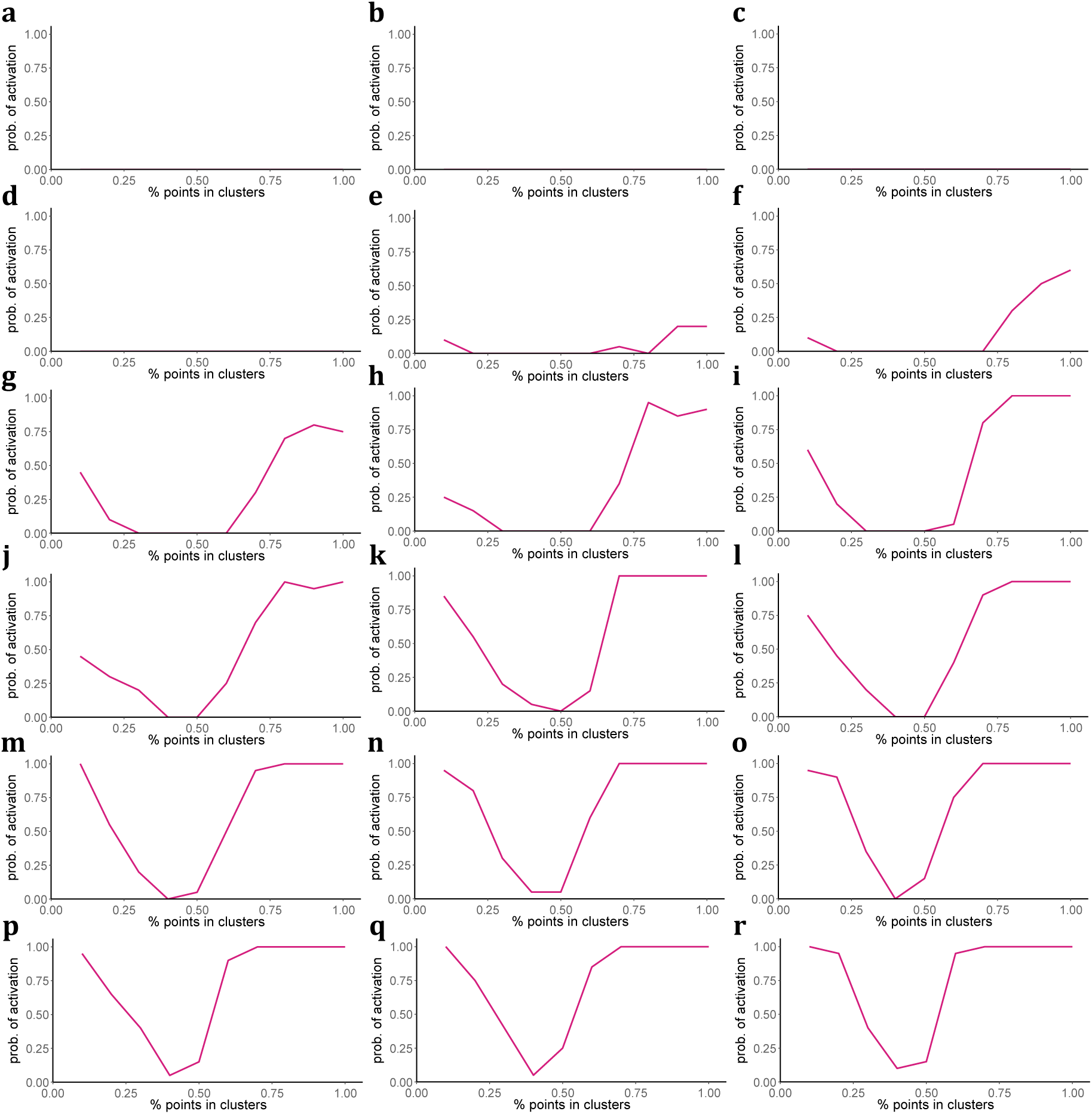
Estimated probability of cell activation versus % points in clusters (with all other parameters fixed) for A/I ratios of **a** 0.25, **b** 0.5, **c** 0.75, **d** 1.0, **e** 1.25, **f** 1.5, **g** 1.75, **h** 2.0, **i** 2.25, **j** 2.5, **k** 2.75, **l** 3.0, **m** 3.25, **n** 3.5, **o** 3.75, **p** 4.0, **q** 4.25 and **r** 4.5.

**Supplementary Figure 4:**
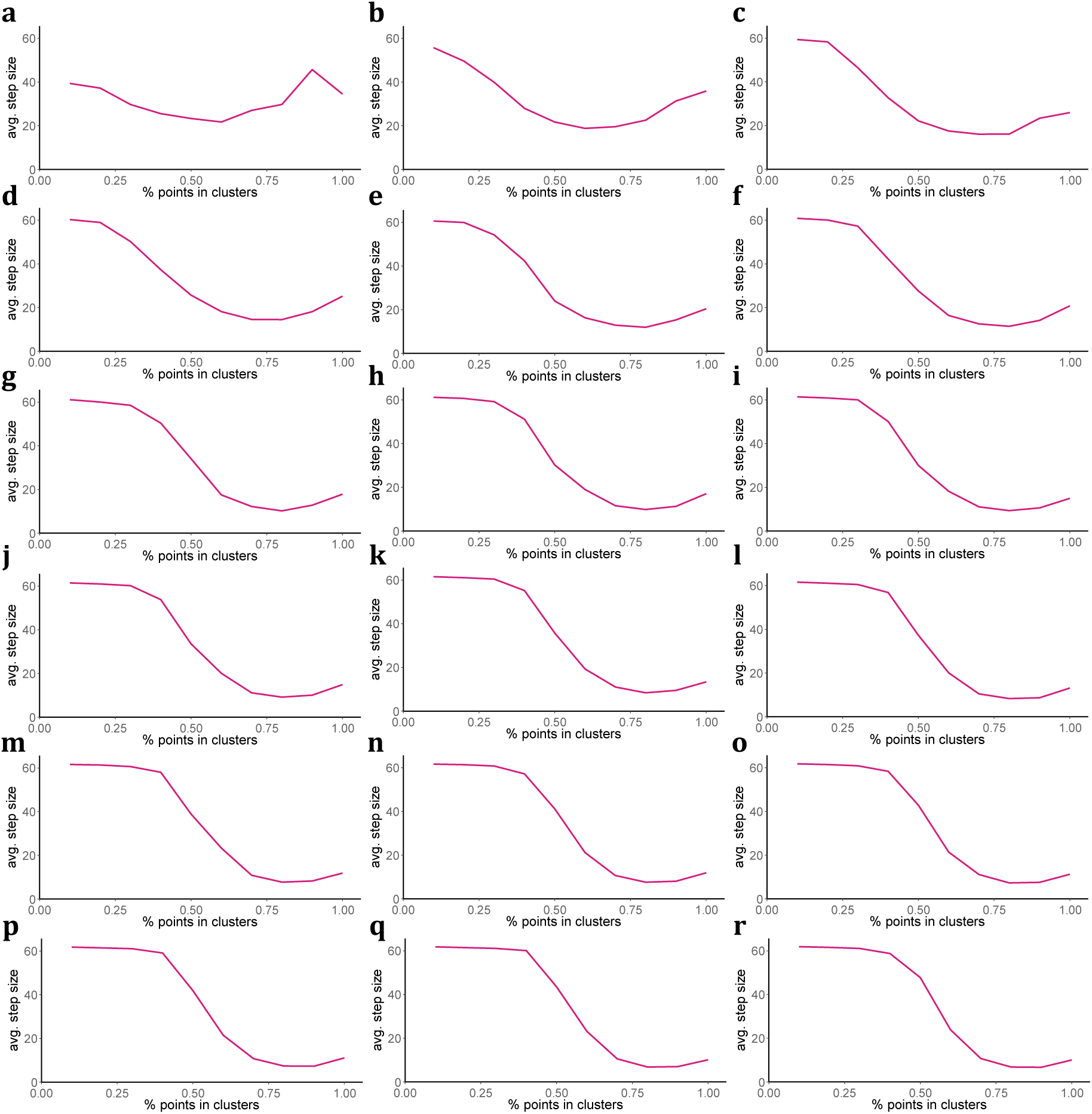
Average step size across all simulations versus % points in clusters (with all other parameters fixed) for A/I ratios of **a** 0.25, **b** 0.5, **c** 0.75, **d** 1.0, **e** 1.25, **f** 1.5, **g** 1.75, **h** 2.0, **i** 2.25, **j** 2.5, **k** 2.75, **l** 3.0, **m** 3.25, **n** 3.5, **o** 3.75, **p** 4.0, **q** 4.25 and **r** 4.5.

## Notes

### Competing Interest Statement

The authors have declared no competing interest.

